# Intraflagellar transport-A deficiency attenuates ADPKD in a renal tubular- and maturation-dependent manner

**DOI:** 10.1101/2020.04.26.061796

**Authors:** Wei Wang, Luciane M. Silva, Henry H. Wang, Matthew A. Kavanaugh, Tana S. Pottorf, Bailey A. Allard, Damon T. Jacobs, Rouchen Dong, Joseph T. Cornelius, Aakriti Chaturvedi, Michele T. Pritchard, Madhulika Sharma, Chad Slawson, Darren P. Wallace, James P. Calvet, Pamela V. Tran

**Author notes:** Correspondence should be addressed to: Pamela V. Tran, Department of Anatomy and Cell Biology and The Jared Grantham Kidney Institute, University of Kansas Medical Center, 3901 Rainbow Blvd., MS #3038, Kansas City, KS 66160.

## Abstract

Primary cilia are sensory organelles built and maintained by intraflagellar transport (IFT) multi-protein complexes. Deletion of different IFT-B genes attenuates polycystic kidney disease (PKD) severity in juvenile and adult Autosomal Dominant (AD) PKD mouse models, while deletion of an IFT-A adaptor, *Tulp3*, attenuates PKD severity in adult mice only. These studies indicate that dysfunction of specific cilia components has potential therapeutic value. To broaden our understanding of cilia dysfunction and its therapeutic potential, here we investigate the impact of global deletion of an IFT-A gene, *Thm1*, in juvenile and adult ADPKD mouse models. Both juvenile and adult models exhibited increased kidney weight:body weight (KW/BW) ratios, renal cysts, inflammation, lengthened renal cilia, and increased levels of the nutrient sensor, O-linked β-N-acetylglucosamine (O-GlcNAc). *Thm1* deletion in juvenile ADPKD mice reduced KW/BW ratios and cortical collecting duct cystogenesis, but increased proximal tubular and glomerular dilations and did not reduce inflammation, cilia lengths, and O-GlcNAc signaling. In contrast, *Thm1* deletion in adult ADPKD mice markedly attenuated renal cystogenesis, inflammation, cilia lengths, and O-GlcNAc. Thus, unlike IFT-B genes, the role of *Thm1* deletion in ADPKD mouse models is development-specific. Unlike an IFT-A adaptor gene, deleting *Thm1* in juvenile ADPKD mice is partially ameliorative. Our studies suggest that different microenvironmental factors found in distinct nephron segments and between developing and mature kidneys modify ciliary homeostasis and ADPKD pathobiology. Further, elevated levels of O-GlcNAc, which regulates cellular metabolism and ciliogenesis, may be a novel feature and critical regulator of certain key ADPKD pathological processes.

## Introduction

Autosomal Dominant Polycystic Kidney Disease (ADPKD) is among the most common, fatal monogenetic diseases, affecting approximately 1:1000 individuals worldwide^1^. ADPKD is characterized by the growth of large fluid-filled renal cysts, which cause injury and fibrosis and can lead to end-stage renal disease by the 6^th^ decade of life. Tolvaptan is the only FDA-approved therapy, but has variable effectiveness and aquaresis side effects^2, 3^. Thus, the need to discover additional underlying disease mechanisms and design new therapeutic strategies continues.

Primary cilia are small, antenna-like sensory organelles that play an important role in ADPKD pathobiology via mechanisms that remain unclear. ADPKD is caused by mutations in *PKD1* or *PKD2*, which encode polycystin 1 (PC1) and polycystin 2 (PC2), respectively^4^. PC1 and PC2 form an ion-channel receptor complex that functions at the primary cilium. While PC1 and PC2 also localize to other subcellular compartments, analyses of human ADPKD primary renal epithelial cells, of mouse models harboring human ADPKD mutations, and of an ethylnitrosourea (ENU)-induced *Pkd2* mouse mutation that causes ciliary exclusion of PC2, indicate that deficiency of polycystins from the cilium is sufficient to cause ADPKD^5–7^.

Primary cilia are synthesized and maintained via intraflagellar transport (IFT), which is the bi-directional transport of protein cargo along a microtubular axoneme. Two multiprotein complexes mediate IFT. The IFT-B complex interacts with the kinesin motor and mediates anterograde IFT, while the IFT-A complex together with cytoplasmic dynein mediates retrograde IFT. IFT-A proteins are also required for ciliary import of membrane and signaling molecules^8–10^. Additionally, an IFT-A adaptor, TULP3, binds to the IFT-A complex and brings in certain G-protein signaling molecules. In mice, deletion of *Ift-A* or *-B* genes or of *Tulp3*, either perinatally or in the embryonic kidney results in renal cystic disease^11–13^. However, these mutants differ from ADPKD models in manifesting generally smaller renal cysts and greater fibrosis relative to cyst size^14, 15^. Additionally, *Ift-A* and *–B* mutants differ in cilia phenotype – often shortened and absent cilia, respectively - and can also show opposing signaling phenotypes, reflecting the differing functions of IFT-A and -B^12, 16–18^. Intriguingly, deletion of *Ift-B* genes, *Kif3a, Ift20* and *Ift88* in juvenile and adult *Pkd1* or *Pkd2* conditional knock-out (cko) mice reduces PKD severity^19–22^, while deletion of *Tulp3* attenuates PKD in adult mice only. Collectively, these studies indicate that a component of cilia dysfunction has potential therapeutic value.

The role of IFT-A deficiency in ADPKD has not been reported. *THM1* (TPR-containing Hedgehog modulator 1; also termed *TTTC21B*) is an ortholog of *Chlamydomonas IFT139*, an IFT-A gene^16^. Causative and modifying mutations in *THM1* have been identified in patients with nephronophthisis, Bardet Biedl syndrome, Meckel syndrome, Jeune syndrome and renal agenesis^14^. Characteristic of IFT-A mutation, deletion of *Thm1* impairs retrograde IFT, causing accumulation of proteins in bulb-like distal tips of shortened primary cilia^16^. *Thm1* loss also impairs cilia entry of membrane-associated proteins, delays and reduces ciliogenesis, and promotes serum-induced cilia loss^23^. In mice, *Thm1* deletion recapitulates many of the clinical manifestations of ciliopathies^16, 24, 25^. Perinatal global deletion of *Thm1* results in renal cystic disease^24^, while deletion of *Thm1* in adult mice does not result in a renal phenotype by 3 months of age, consistent with the developmental time-frame that determines whether loss of a cystogenic gene will cause rapid- or slow-progressing renal cystic disease^26^. To expand on the role of ciliary dysfunction in ADPKD, here we investigate the role of *Thm1*/ IFT-A deficiency in juvenile and adult ADPKD mouse models.

## Results

### Perinatal deletion of *Thm1* in *Pkd2* conditional knock-out mice reduces cortical collecting duct cystogenesis, but does not improve kidney function

To examine the effect of IFT-A deficiency in a rapidly progressing ADPKD mouse model, we deleted *Thm1* together with *Pkd2* at postnatal day (P) 0, and examined the renal phenotypes of control, *Thm1* conditional knock-out (cko), *Pkd2* cko and *Pkd2;Thm1* double knock-out (dko) mice at P21. At this early stage, *Thm1* cko mice on a pure C57BL6/J background showed kidneys with some cortical tubular dilations, reduced kidney weight/body weight (KW/BW) ratios, and elevated blood urea nitrogen (BUN) levels (Figures 1A-1C). *Pkd2* cko mice showed cysts in both renal cortex and medulla, as well as increased KW/BW ratios and BUN levels. Relative to *Pkd2* cko mice, *Pkd2;Thm1* dko mice showed decreased cystogenesis specifically in the cortex (Figures 1D-1F), reduced KW/BW ratios, but similar BUN levels, indicating kidney dysfunction was not improved.

**Figure 1.**
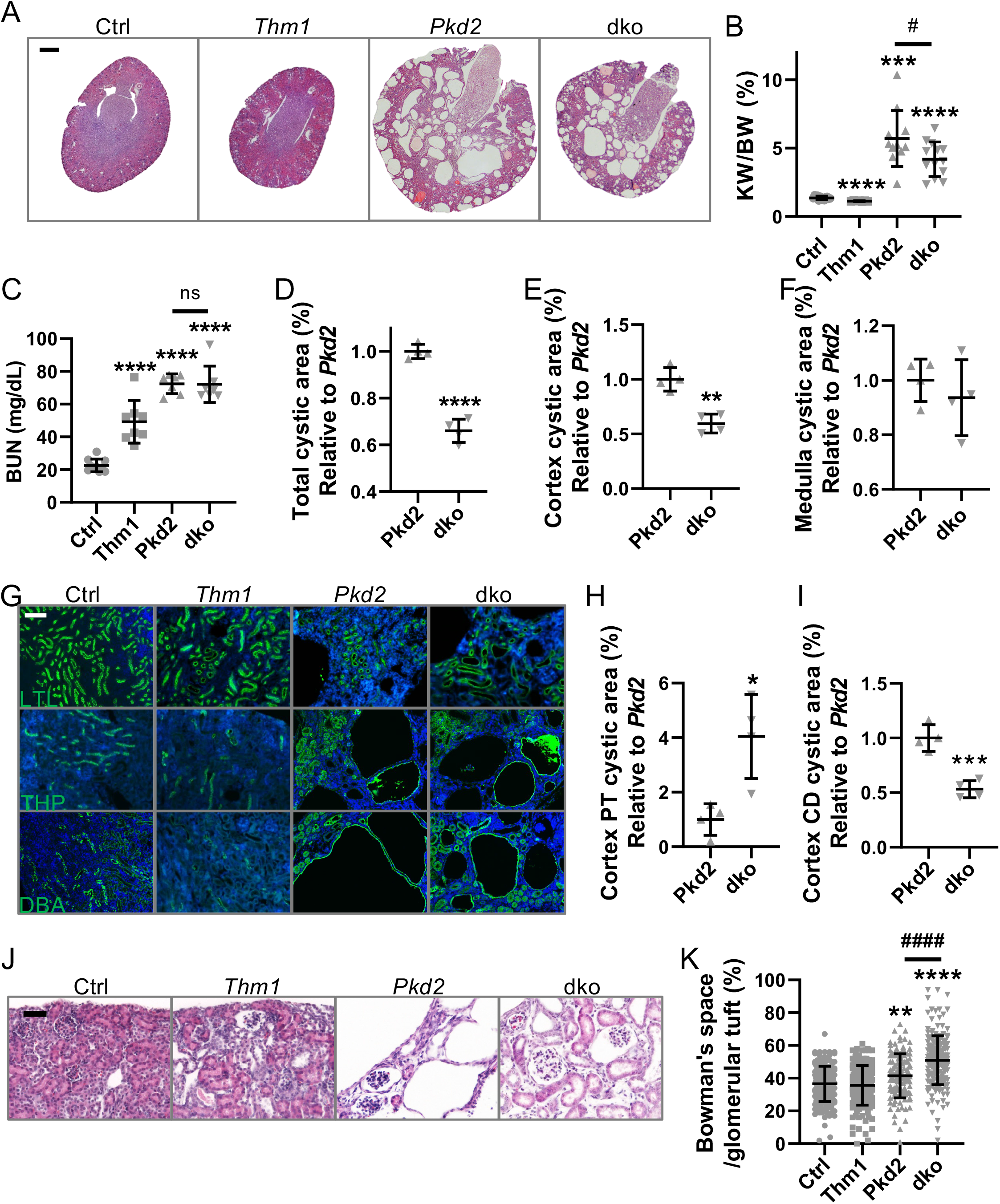
*Thm1* deletion in juvenile *Pkd2* conditional knock-out mice reduces cortical cystogenesis. (A) H&E staining of P21 kidney sections. Scale bar - 500μm. (B) KW/BW ratios (C) BUN levels (D) Percent cystic index of whole kidney; (E) of cortex; (F) and of medulla. *Pkd2* cko values are set at 1. Bars represent mean ± SD. Statistical significance was determined by Brown-Forsythe and Welch ANOVA tests followed by Dunnett’s T3 multiple comparisons test in (B), and ordinary one-way ANOVA test followed by Sidak’s multiple comparisons test in (C). ***p<0.001, ****p<0.0001, compared to Ctrl. In (B) and (C), unpaired t-test was used to determine the difference between *Pkd2* and dko groups. ^##^p<0.05. In (D), (E) and (F), statistical significance was determined by unpaired two-tailed t-test. **p<0.01, ****p<0.0001, compared to Ctrl. Each data point represents a mouse. (G) Staining of kidney cortex with LTL, THP and DBA. Scale bar - 100μm (H) Quantification of LTL+ dilations and (I) DBA+ cystic index in renal cortex. *Pkd2* cko values are set at 1. Bars represent mean ± SD. Statistical significance was determined by unpaired two-tailed t-test. *p<0.05, ***p<0.001, compared to Ctrl. (J) H&E staining. Scale bar - 50μm (K) Area of Bowman’s space/area of Bowman’s capsule. Bars represent mean ± SD. Statistical significance was determined by Kruskal-Wallis test followed by Dunn’s multiple comparisons test. ^####^p<0.0001; ^**\*\***^p<0.01 compared to Ctrl; ^**\*\*\*\***^p<0.0001 compared to Ctrl

We examined the tubular origin of the renal cortical dilations and cysts. *Thm1* cko renal cortical dilations were mostly LTL+, indicating proximal tubules, while fewer were THP+ or DBA+, marking loop of Henle and collecting duct, respectively (Figure 1G). *Pkd2* cko renal cortices showed LTL+ dilations, THP+ cysts, and multiple, large DBA+ cysts. Relative to *Thm1* cko and *Pkd2* cko kidneys, *Pkd2; Thm1* dko cortices showed greater LTL+ dilations (Figures 1G, 1H). Relative to *Pkd2* cko kidneys, *Pkd2; Thm1* dko cortices showed similar THP+ cystogenesis (Figures 1G, S1A), but decreased DBA+ cysts (Figures 1G, 1I). Glomerular dilations were also present in *Pkd2* cko kidneys, and increased in *Pkd2;Thm1* dko kidneys (Figures 1J, 1K). These data reveal nephron segment-specific effects of *Thm1* deletion on a *Pkd2* cko background.

### *Pkd2* deletion increases cilia length on renal epithelia

In control kidneys, ciliary axonemes marked by acetylated α-tubulin were 3.0μm and 2.1μm for LTL+ and DBA+ cortical tubular epithelial cells, respectively (Figure 2). We also noted qualitative differences between LTL+ and DBA+ primary cilia, with the former cilia being thinner and longer, and the latter being thicker and more rod-like. Cilia lengths of LTL+ and DBA+ cells were increased in *Pkd2* cko cortices. However, relative to *Pkd2* cko cells, cilia lengths of *Pkd2;Thm1* dko LTL+ cells were further increased, but similar for DBA+ cells. These differences reveal tubular-specific effects on cilia phenotype.

**Figure 2.**
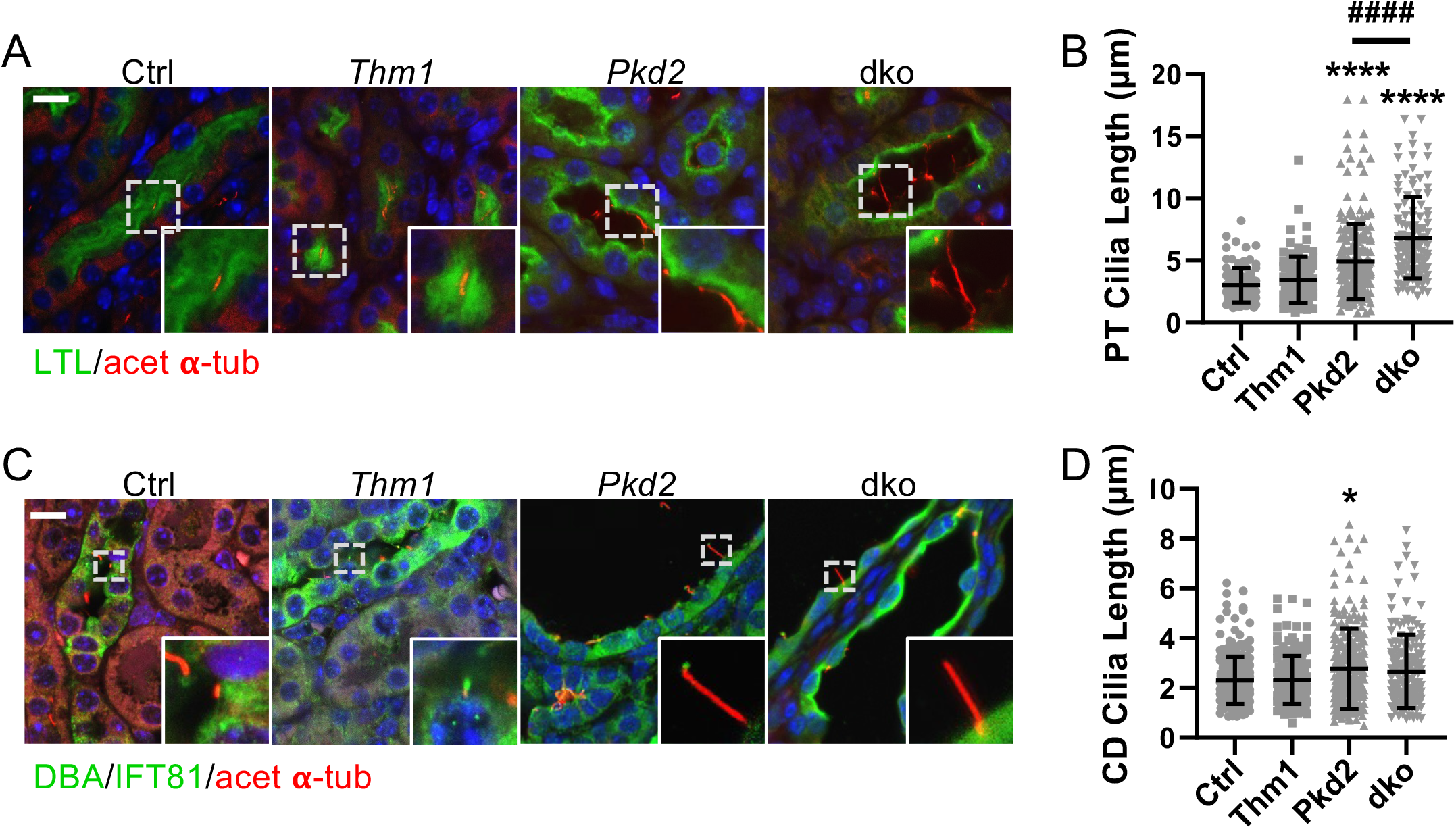
Perinatal *Pkd2* deletion increases renal epithelial primary cilia lengths. (A) Images of kidney cortex immunostained for acetylated α-tubulin (red) together with LTL (green). Scale bar - 10μm. (F) Quantification of cilia length of LTL+ cells. (G) Immunostaining of kidney cortex for acetylated α-tubulin and IFT81 together with DBA. Scale bar - 10μm. n=3 mice/genotype. Quantification of cilia length of cortical DBA+ cells. Bars represent mean ± SD. Statistical significance was determined by Kruskal-Wallis test followed by Dunn’s multiple comparisons test. *p<0.05, ****p<0.0001, compared to Ctrl; ^####^p<0.0001.

ARL13B is a ciliary membrane protein essential for ciliogenesis^27^, and in cultured *Thm1*-null mouse embryonic fibroblasts, ARL13B in cilia is reduced^23^. We therefore examined whether ARL13B may have a role in the renal ciliary phenotypes. However, in DBA+ cortical renal epithelial cells, ARL13B intensities were similar across the mutant genotypes (Figure S2).

We examined whether *Thm1* expression differed in proximal tubules versus cortical collecting duct. However, X-gal staining on kidneys of mice harboring a *Thm1-lacZ* allele showed relatively ubiquitous expression in the cortex and present in both LTL+ and cortical DBA+ tubules (Figure S3).

### *Thm1* deletion increases inflammation

To further evaluate the effect of *Thm1* deletion on disease severity, we examined proliferation. Nuclear staining of proliferating cell nuclear antigen (PCNA) of LTL+ tubules was similar across control, *Thm1* cko and *Pkd2* cko, but slightly elevated in *Pkd2;Thm1* dko kidneys (Figures 3A, 3B). In contrast, PCNA+ nuclei were increased in *Pkd2* cko and *Pkd2;Thm1* dko DBA+ non-cystic tubules, and further increased in *Pkd2* cko and *Pkd2;Thm1* dko DBA+ cysts (Figure 3C). These data support that increased proliferation is an early driver of ADPKD renal cystogenesis.

**Figure 3.**
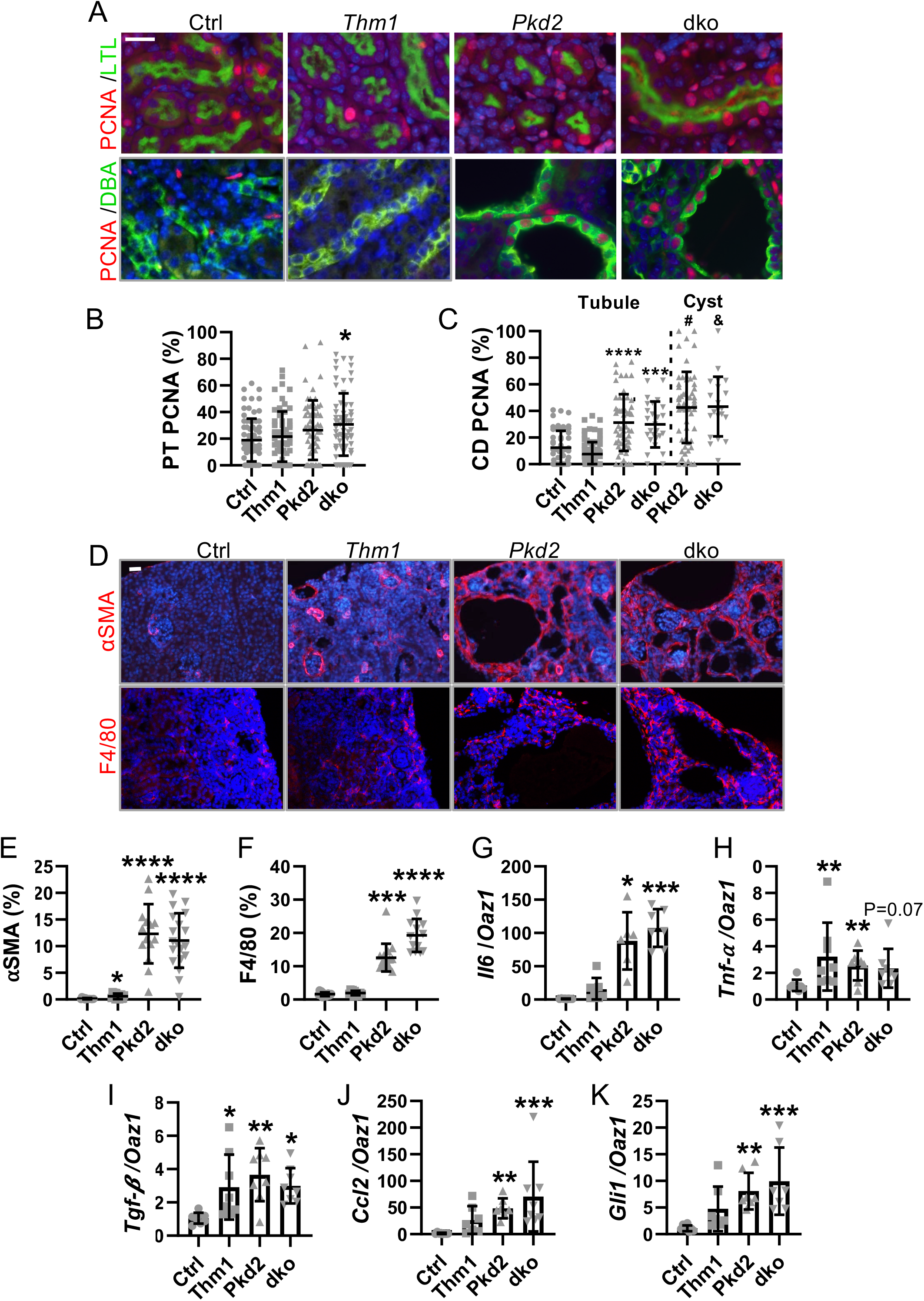
Perinatal *Thm1* deletion increases inflammation. (A) Images of kidney cortex immunostained for PCNA (red) together with LTL or DBA (green). Scale bar - 10μm. (B) Percent PCNA+ cells per LTL+ and (C) DBA+ tubule. Each dot represents a tubule or a cyst from n=3 mice/genotype. Bars represent mean ± SD. Statistical significance was determined by Kruskal-Wallis test followed by Dunn’s multiple comparisons test. *p<0.05, ***p<0.001, ****p<0.0001, compared to Ctrl. Unpaired t-test was used to determine the difference between non-cystic tubules and cysts. ^#^P<0.05 compared to *Pkd2* cko non-cystic tubule; ^&^P<0.05 compared to dko non-cystic tubule. (D) Immunostaining for αSMA (red) and F4/80 (red). Scale bar - 50μm. (E) Quantification of αSMA and (F) F4/80 staining. Each dot represents percentage of staining of a 20X or 40X image (4-5 images/mouse) from n=3 mice/genotype. (G, H, I, J, K) qPCR. Each dot represents a mouse. In (E) and (G), statistical significance was determined by Brown-Forsythe and Welch ANOVA tests followed by Dunnett’s T3 multiple comparisons test. In (F), (H), (J) and (K), statistical significance was determined by Kruskal-Wallis test followed by Dunn’s multiple comparisons test. In (I), statistical significance was determined by ordinary one-way ANOVA followed by Sidak’s multiple comparisons test. *p<0.05, **p<0.01, ***p<0.001, ****p<0.0001, compared to Ctrl. Bars represent mean ± SD.

Cyst growth compresses surrounding parenchyma, leading to injury, inflammation, and fibrosis. We immunostained kidney sections for alpha smooth muscle actin (αSMA) and F4/80 to examine the presence of myofibroblasts and macrophages, respectively, which contribute to pro-inflammatory and pro-fibrotic processes. While control kidneys showed αSMA around blood vessels and a few F4/80+ cells, *Thm1* cko kidneys showed increased αSMA around glomeruli and tubular dilations, and *Pkd2* cko and *Pkd2;Thm1* dko kidney showed even greater αSMA+ and increased F4/80+ labelling surrounding glomeruli, tubular dilations and cysts (Figure 3D-3F). Additionally, transcripts of inflammatory molecules, *Tnfα* and *Tgfβ*, were increased in *Thm1* cko renal extracts, while those of *Il6, Tnfα, Tgfβ, Ccl2* and *Gli1* were increased in *Pkd2* cko and *Pkd2;Thm1* dko kidney extracts (Figures 3G-3K). Thus, deletion of *Thm1* alone increases inflammatory processes, and its deletion on a *Pkd2* cko background results in similar or slightly increased ADPKD inflammation at P21.

### Perinatal deletion of *Thm1* increases ERK and STAT3 signaling

Increased ERK and STAT3 signaling promote disease progression in ADPKD^28–31^. ERK activation promotes cell proliferation^30, 32^ and also acts upstream of mTOR and AMPK pathways regulating cellular metabolism^33^. STAT3 activation can have proliferative and inflammatory roles^31, 34, 35^. At P21, P-ERK did not localize with LTL, but localized with DBA (Figure 4A). While a similar number of P-ERK+ tubules was observed across control and the mutant genotypes (Figure 4B), P-ERK intensity was increased in dilated tubules of *Thm1* cko mice, and further increased in cyst-lining cells of *Pkd2* cko and *Pkd2;Thm1* dko mice (Figure 4C). These data support that ERK activation is a driver of renal cyst growth. Using immunohistochemistry, P-STAT3 was revealed to be increased in epithelial cells lining cortical dilations in *Thm1* cko kidneys and to be increased further in cyst-lining cells of *Pkd2* cko and *Pkd2;Thm1* dko kidneys, both in the cortex and medulla (Figure 4E). Western blot analyses also reveal increased ERK and STAT signaling in *Pkd2* cko and *Pkd2;Thm1* dko renal extracts (Figures 4F, 4G). Thus, *Thm1* loss as well as *Pkd2* cystic disease increase ERK and STAT3 signaling.

**Figure 4.**
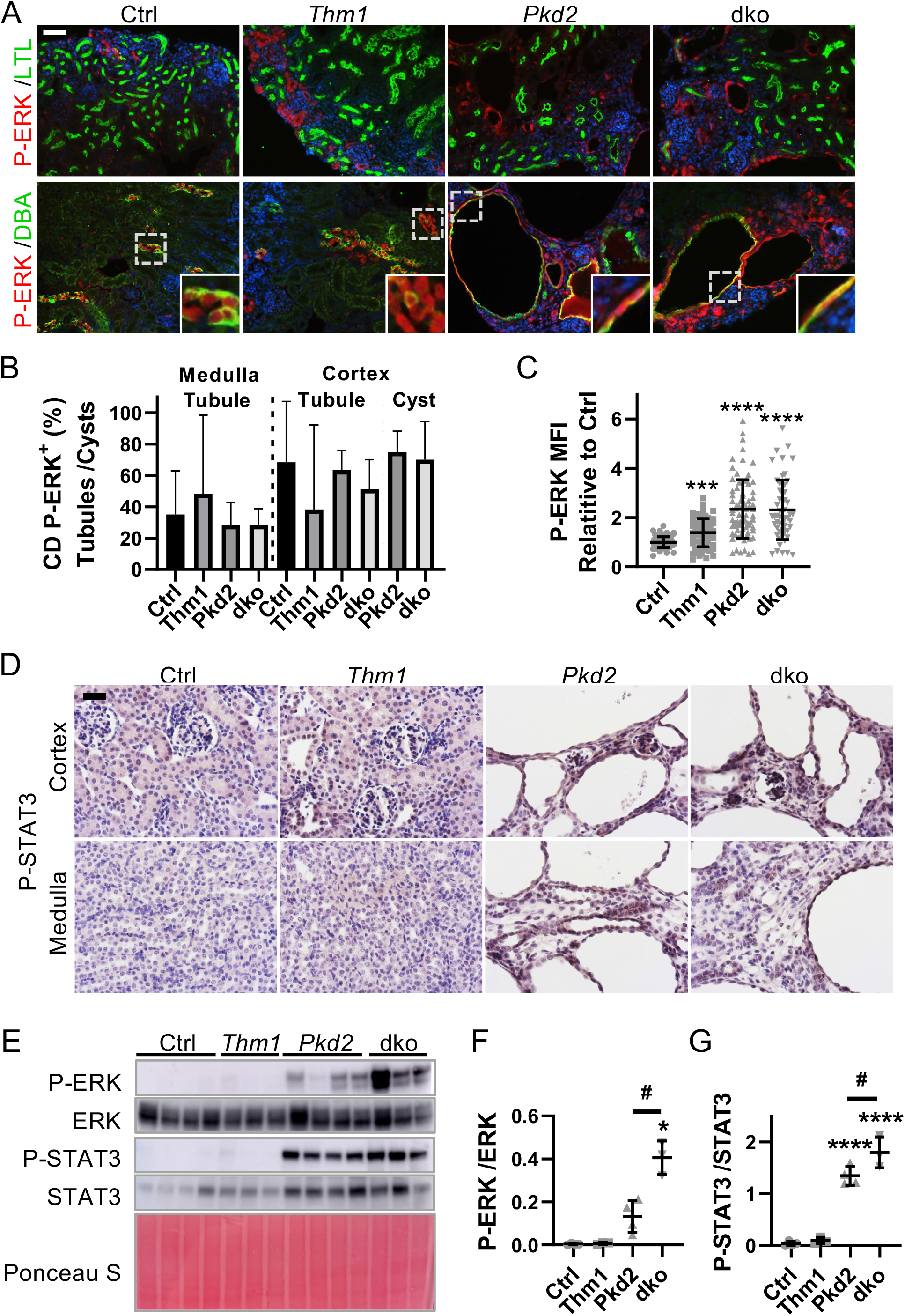
Perinatal *Thm1* deletion increases ERK and STAT3 activation. (A) Images of kidney cortex immunostained for P-ERK together with LTL or DBA. Scale bar - 50μm. (B) Quantification of P-ERK+ tubules or cysts and (C) P+ ERK intensity. Each dot represents a tubule or cyst from n=3 mice/genotype. Statistical significance was determined by Kruskal-Wallis test followed by Dunn’s multiple comparisons test. ***p<0.001, ****p<0.0001, compared to Ctrl. (D) Immunohistochemistry for P-STAT3. n=3 mice/genotype. Scale bar - 25μm. (E) Western blot for STAT3 and ERK signaling, (F) Quantification of P-STAT3/STAT3 and (G) of P-ERK/ERK ratios. Statistical significance was determined by ordinary one-way ANOVA test followed by Sidak’s multiple comparisons test or Brown-Forsythe and Welch ANOVA tests followed by Dunnett’s T3 multiple comparisons test, respectively. *p<0.05, ****p<0.0001, compared to Ctrl. ^#^p<0.05.

### O-GlcNAc is increased in dilated tubules and cysts

Altered cellular metabolism has emerged as another component of ADPKD pathobiology^33, 36–38^. One of these alterations includes the Warburg effect^33^, whereby cells preferentially convert the product of glycolysis, pyruvate, into lactate, even in the presence of oxygen, at the expense of pyruvate entering mitochondria to enable oxidative phosphorylation. The nutrient sensor, O-linked β-N-acetylglucosamine (O-GlcNAc) regulates the balance between glycolysis and oxidative phosphorylation^39^, as well as ciliary homeostasis^40, 41^, both of which are altered in ADPKD^33, 42^. We therefore hypothesized that O-GlcNAc signaling is misregulated in PKD. Relative to control, *Thm1* cko kidneys showed increased O-GlcNAc expression in nuclei of cells lining dilations in the cortex, while *Pkd2* cko and *Pkd2;Thm1* dko kidneys showed increased OGlcNAc staining in cyst-lining cells in both the cortex and medulla (Figures 5A, S4). O-GlcNAC levels are regulated by two enzymes, O-GlcNAc transferase (OGT) and O-GlcNAcase (OGA), which transfers and removes the O-GlcNAc moiety on protein substrates, respectively. In *Thm1* cko kidneys, OGT was increased in the cytoplasm and nuclei of cells lining cortical dilations. In *Pkd2* cko and in *Pkd2;Thm1* dko mice, intense OGT staining was present in cyst-lining cells in the cortex and medulla. In control and *Thm1* cko kidneys, light OGA staining was present in cytoplasm of cortical and medullary tubules. In *Pkd2* cko and in *Pkd2;Thm1* dko mice, OGA was increased in the nuclei and cytoplasm of cyst-lining cells in both the cortex and medulla. Western blot analyses showed increased O-GlcNAc levels in mutant renal extracts, with highest levels in *Pkd2* cko renal extracts (Figures 5B, 5C). These data suggest that increased O-GlcNAc signaling is a feature of ADPKD.

**Figure 5.**
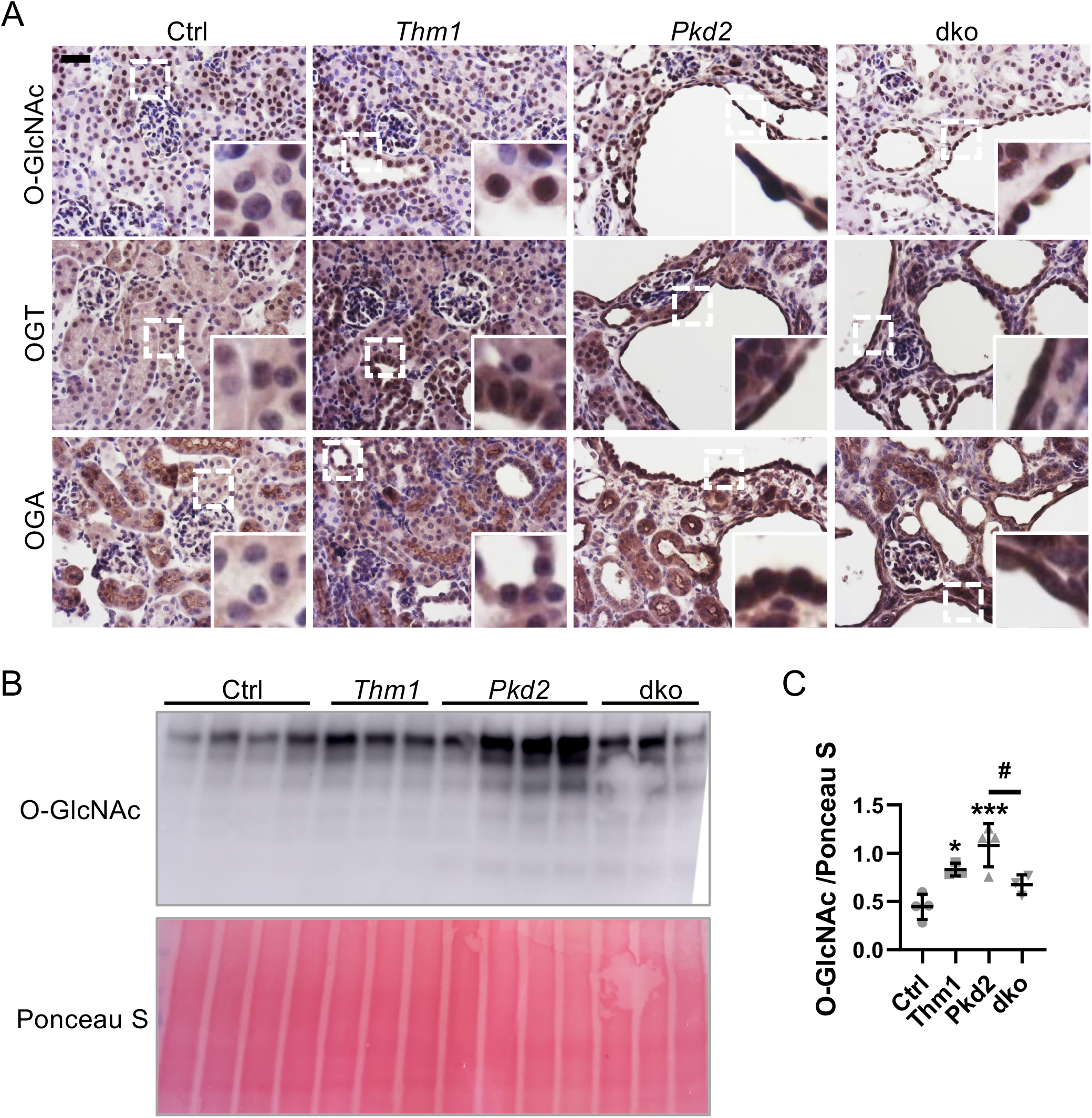
O-GlcNAc is increased in cells lining dilations and cysts in juvenile mice. (A) Images of kidney cortex following immunohistochemistry for O-GlcNAc, OGT and OGA. Scale bar - 25μm. n=3 mice/genotype. (B) Western blot of O-GlcNAc and (C) quantification. Statistical significance was determined by ordinary one-way ANOVA test followed by Sidak’s multiple comparisons test. *p<0.05, ***p<0.001, compared to Ctrl. ^#^p<0.05.

### Deletion of *Thm1* in adult *Pkd2* or *Pkd1* conditional knock-out mice markedly attenuates ADPKD renal cystogenesis

We next examined the role of IFT-A deficiency in slowly progressive adult ADPKD mouse models. We deleted *Thm1* together with *Pkd1* at P35 and examined renal phenotypes at 6 months of age. *Thm1* cko mice showed normal kidney morphology and BUN levels (Figure S5A). In *Pkd1* cko mice, renal cysts were mostly in the cortex, with the largest and most abundant cysts being DBA+, fewer cysts being THP+ and dilations being LTL+ (Figure 6A). Notably, all these features were reduced in *Pkd1;Thm1* dko kidneys. Similarly, KW/BW ratios and BUN levels were elevated in *Pkd1* cko mice, and corrected in *Pkd1;Thm1* dko mice (Figures 6B, 6C).

**Figure 6.**
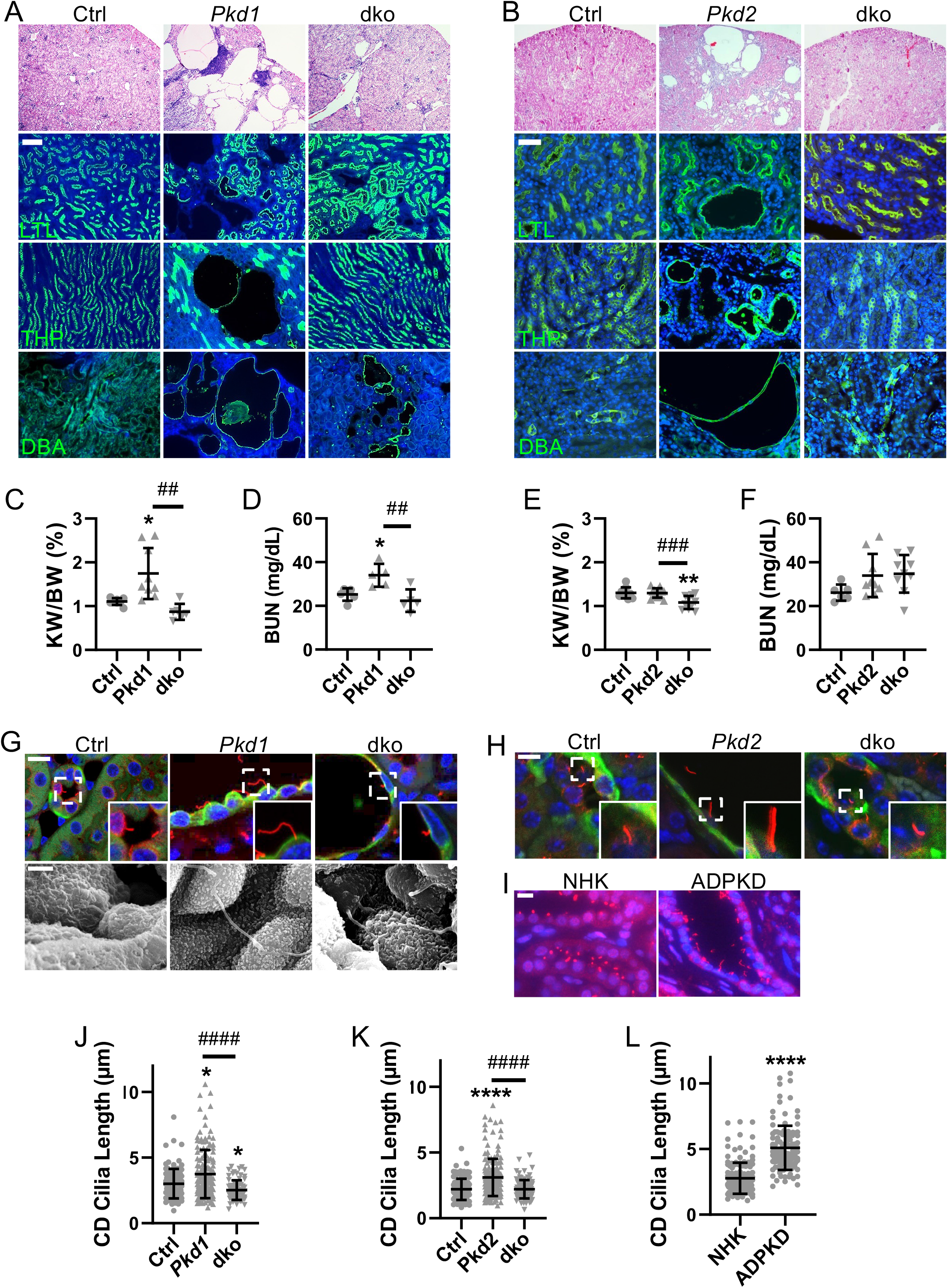
*Thm1* deletion in adult ADPKD mouse models attenuates renal cystogenesis and cilia lengthening. (A) Histology, lectin staining and immunostaining for *Pkd1* cko mice and (B) for *Pkd2* cko mice. Scale bars - 100μm and 50μm, respectively. (C) KW/BW ratios and (D) BUN levels for *Pkd1* cko mice and (E, F) for *Pkd2* cko mice. Bars represent mean ± SD. Each dot represents an animal. In (C), statistical significance was determined by Brown-Forsythe and Welch ANOVA tests followed by Dunnett’s T3 multiple comparisons test. In (D) and (E), statistical significance was determined by one-way ANOVA followed by Tukey’s test. In (F), statistical significance was determined by Kruskal-Wallis test followed by Dunn’s multiple comparisons test. *p<0.05, **p<0.01, compared to Ctrl; ^##^p<0.01; ^###^p<0.001. (G) Images of *Pkd1* cko kidney cortex immunostained for acetylated α-tubulin together with DBA. Scale bar - 10μm. Scanning electron micrographs of primary cilia. Scale bar – 1.5μm. n=3 mice/genotype. (H) Images of *Pkd2* cko kidney cortex immunostained for acetylated α-tubulin together with DBA. Scale bar - 10μm. n=3/genotype. (I) Immunostaining for ARL13B of normal human kidney (NHK) and ADPKD renal sections. Scale bar - 10μm. n=3 for NHK and ADPKD. (J) Quantification of renal cilia lengths of *Pkd1* cko mice, (K) *Pkd2* cko mice, and (L) ADPKD tissue. Cilia lengths were quantified from immunofluorescence experiments of (G, H, I). Significance was determined using Kruskal-Wallis test followed by Dunn’s multiple comparisons test (J, K) or by Mann Whitney test (L). Bars represent mean ± SD. ^####^p<0.0001; *p<0.05, ****p<0.0001, compared to Ctrl.

Since *PKD2* mutations result in less severe PKD, we deleted *Thm1* together with *Pkd2* at P28 and examined renal phenotypes at 6 months of age. *Thm1* cko kidney morphology and BUN levels resembled control (Figure S5B). *Pkd2* cko mice show renal cysts mostly in the cortex, with the largest cysts being DBA+, and smaller cysts being LTL+ or THP+ (Figure 6D). In contrast, in *Pkd2;Thm1* dko mice, the *Pkd2* cko cystic phenotype is largely corrected morphologically. In *Pkd2* cko mice, KW/BW ratios were similar to control and BUN levels showed a trend toward a slight elevation, reflecting the mild disease induced in adulthood. In *Pkd2;Thm1* dko mice, KW/BW ratios were slightly reduced, due to increased body weight caused by global deletion of *Thm1*^*25*^, and BUN levels showed a slightly decreasing pattern relative to *Pkd2* cko mice.

### Deletion of *Thm1* in adult ADPKD mouse models reduces cilia lengths of cortical collecting duct renal epithelia

Similar to juvenile ADPKD models, cilia lengths of *Pkd1* cko and *Pkd2* cko DBA+ adult renal epithelial cells were increased (Figures 6G, 6H, 6J, 6K). However, in contrast to juvenile models, cilia lengths of *Pkd1;Thm1* and *Pkd2;Thm1* dko DBA+ epithelia were reduced relative to those of *Pkd1* cko and *Pkd2* cko epithelia and similar to control. Additionally, human ADPKD sections had longer renal epithelial cilia than normal human kidney (NHK) sections (Figures 6I, 6L), supporting that increased cilia length is also a feature of the human disease^22^.

### Deletion of *Thm1* in adult *Pkd1* conditional knock-out mice reduces proliferation, inflammation, P-ERK, P-STAT3, and O-GlcNAc

We examined the extent of ADPKD attenuation by *Thm1* deletion. *Pkd1* cko kidneys showed increased PCNA in cyst-lining cells, as well as increased αSMA and F4/80 around cysts and glomeruli, which were markedly reduced in *Pkd1;Thm1* dko kidneys (Figures 7A-D). Transcripts of pro-inflammatory molecules, *Ccl2, Il6, Tnfα, Tgfβ, Ccl2* and *Gli1*, were elevated in *Pkd1* cko kidneys, but reduced in *Pkd1;Thm1* dko kidneys (Figure 7E). Similarly, P-ERK and P-STAT3 immunostaining was increased in *Pkd1* cko kidneys and reduced in *Pkd1;Thm1* dko kidneys (Figures 7J, 7K). Thus, proliferative and pro-inflammatory pathways are attenuated in late-onset ADPKD by deletion of *Thm1*.

**Figure 7.**
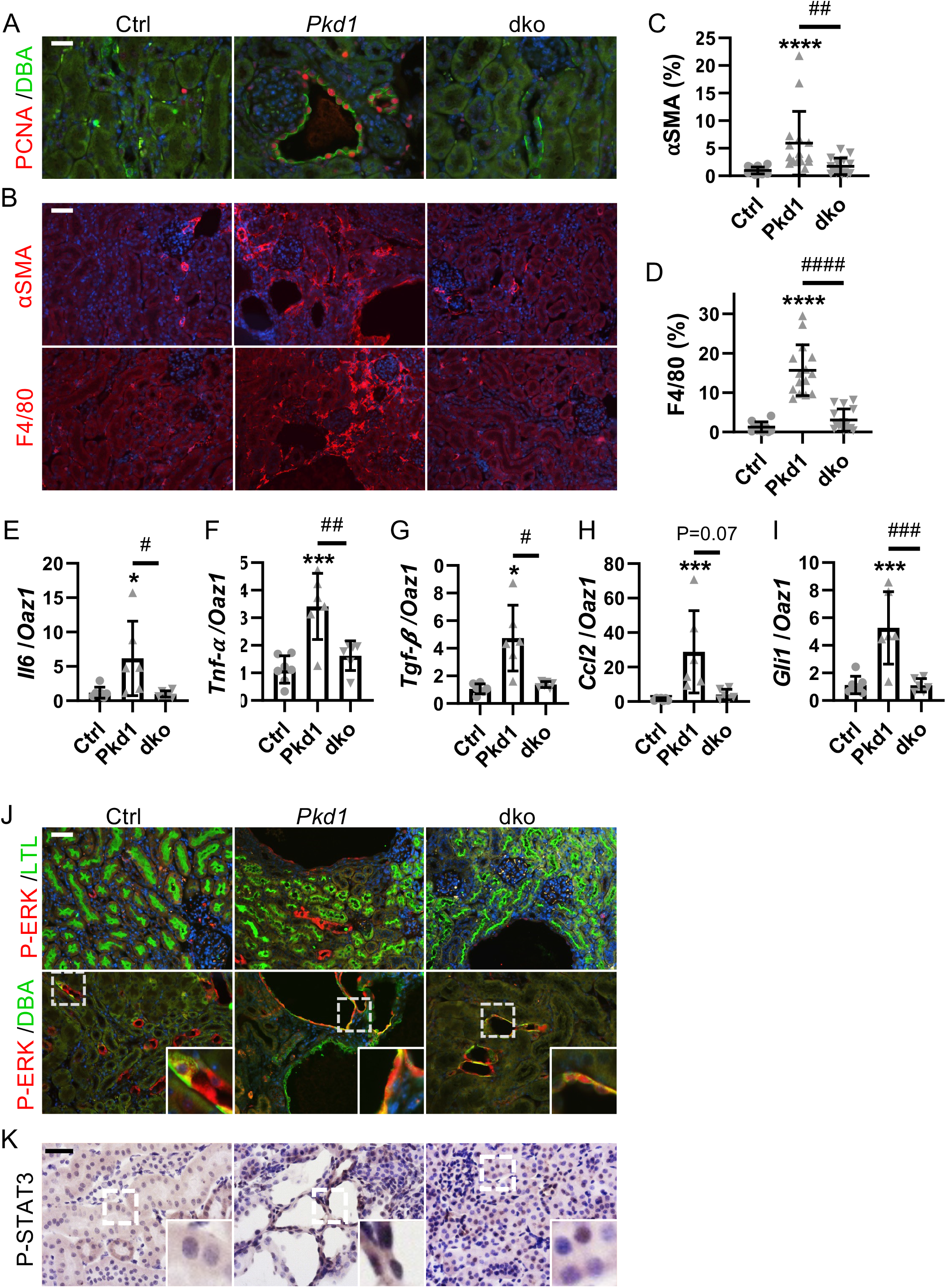
*Thm1* deletion in adult *Pkd1* conditional knock-out mice attenuates proliferation and inflammation. (A) Images of kidney cortex immunostained for PCNA (red) together with DBA (green). Scale bar - 100μm. (B) Immunostaining for αSMA (red) and F4/80 (red). Scale bar - 50μm. (C) Quantification of αSMA and (D) F4/80 staining. Each dot represents percentage of staining of a 20X or 40X image (5 images/mouse) from n=3 mice/genotype. (E, F, G, H, I) qPCR. Each dot represents a mouse. In (C), (D), (E) and (H), statistical significance was determined by Kruskal-Wallis test followed by Dunn’s multiple comparisons test. In (F) and (I), statistical significance was determined by ordinary one-way ANOVA test followed by Tukey’s multiple comparisons test. In (G), statistical significance was determined by Brown-Forsythe and Welch ANOVA tests followed by Dunnett’s T3 multiple comparisons test. *p<0.05, ***p<0.001, compared to Ctrl; ^#^p<0.05, ^##^p<0.01, ^###^p<0.001. Bars represent mean ± SD. (J) Immunostaining for P-ERK (red) together with LTL (green) or DBA (green). Scale bar - 50μm. (K) Immunohistochemistry for P-STAT3. n=3 mice/genotype. Scale bar – 20μm.

Finally, we examined O-GlcNAcylation. In *Pkd1* cko kidneys, increased O-GlcNAc was present in cyst-lining epithelia of the cortex and in proteinaceous substances within some cysts, and in tubules of the medulla (Figures 8A, S6). OGT and OGA were also increased in cyst-lining cells of the cortex and in some tubules of the medulla. In *Pkd1;Thm1* dko kidneys, O-GlcNAc, OGT and OGA staining were reduced. Western blot showed increased O-GlcNAc levels in *Pkd1* cko renal extracts (Figure 8B), and levels were reduced in *Pkd1;Thm1* dko extracts. These data suggest that increased O-GlcNAcylation is a feature also of slowly progressive ADPKD mouse models. Further, deletion of *Thm1* in adult ADPKD mouse models attenuates perturbation of this metabolic regulator.

**Figure 8.**
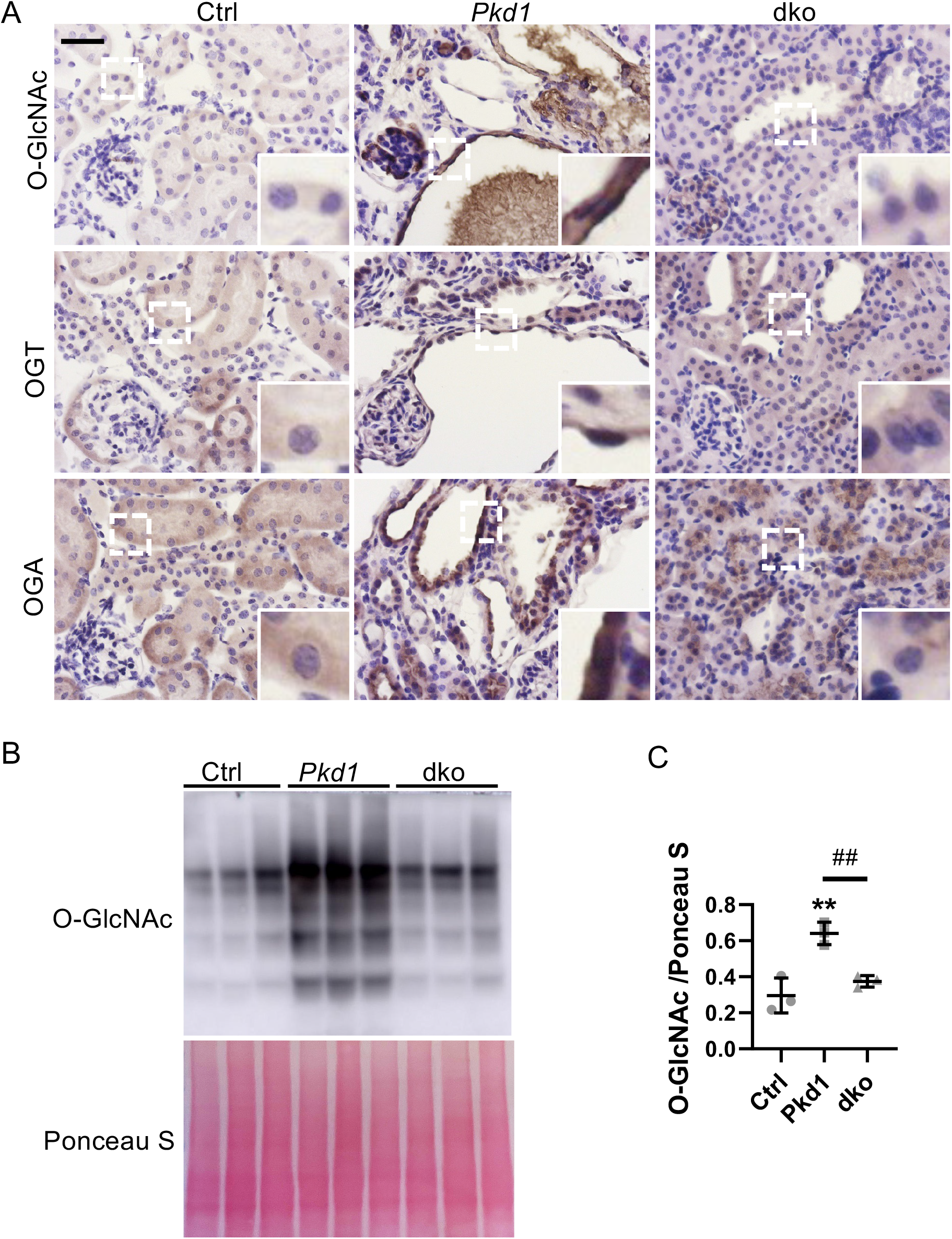
*Thm1* deletion in adult *Pkd1* conditional knock-out mice attenuates increased O-GlcNAc. (A) Images of kidney cortex following immunohistochemistry for O-GlcNAc, OGT and OGA. Scale bar – 20μm. n=3 mice/genotype. (B) Western blot for O-GlcNAc and (C) quantification. Statistical significance was determined by ordinary one-way ANOVA test followed by Tukey’s multiple comparisons test. **p<0.01, compared to Ctrl. ^##^p<0.01.

## Discussion

To expand on the ciliary components whose dysfunction in ADPKD mouse models can attenuate disease severity^19, 21, 43^, we examined the role of deletion of IFT-A gene, *Thm1*. Distinct from IFT-B, deletion of *Thm1* does not improve kidney function in a juvenile ADPKD mouse model, but lessens most disease aspects in an adult model (Figure 9). Distinct from IFT-A adaptor, *Tulp3*, deletion of *Thm1* in a juvenile ADPKD mouse model had partially ameliorative effects, reducing KW/BW ratios and cortical collecting duct cystogenesis. Our data also reveal that the effects of *Thm1* deletion in juvenile ADPKD mice are nephron-specific, with no effect on loop of Henle-derived cysts, and exacerbation of proximal tubular and glomerular dilations. This suggests varying microenvironments between nephron segments modify cilia dysfunction. The differential effects of *Tulp3* or *Thm1* deletion in juvenile versus adult ADPKD mouse models further suggest distinct microenvironments between developing versus mature kidneys that influence not just renal cystogenesis, but also inflammation.

**Figure 9.**
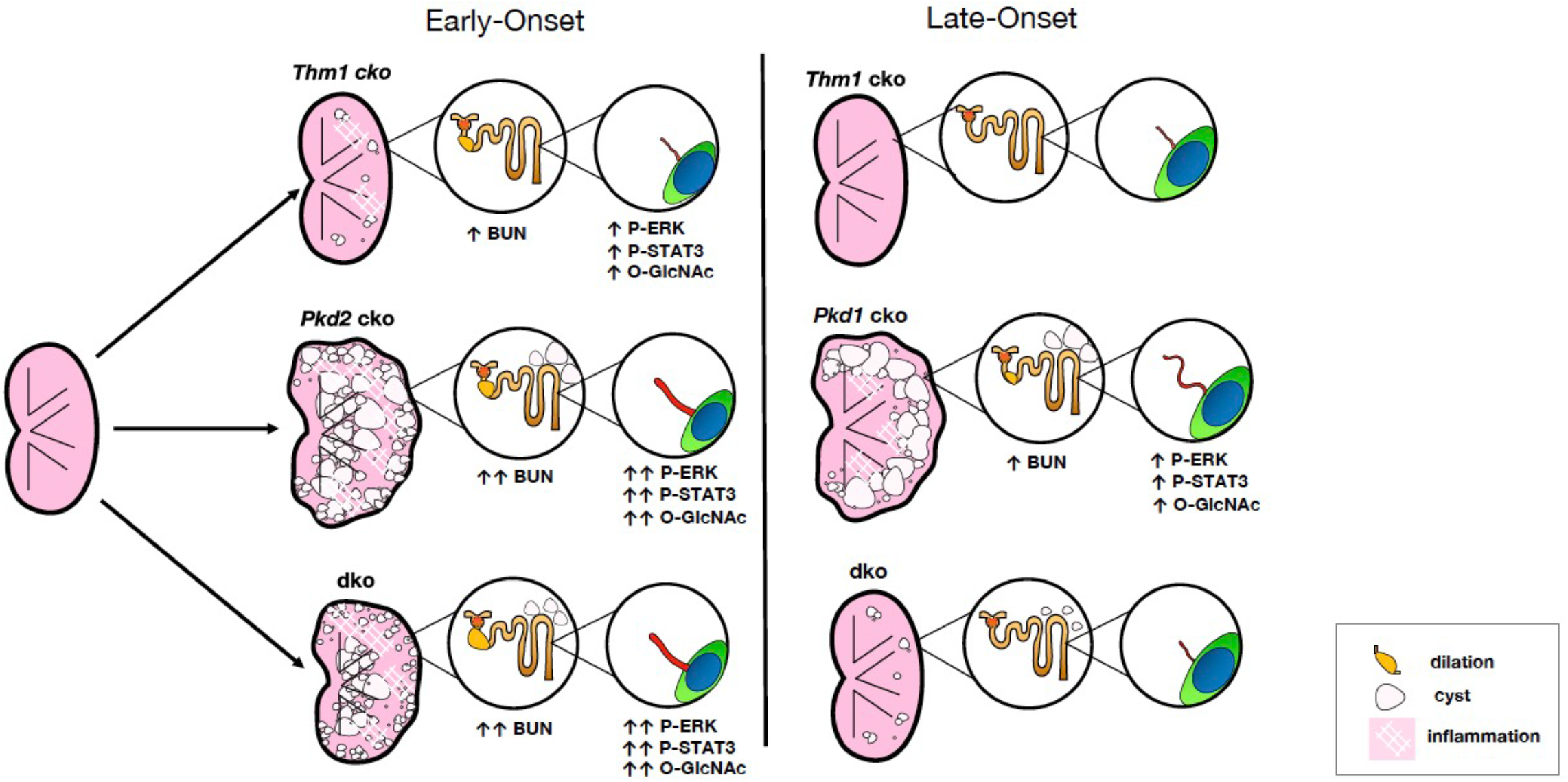
Model of role of IFT-A deficiency in ADPKD. In an early-onset, juvenile ADPKD mouse model, deletion of *Thm1* causes nephron segment-specific effects, attenuating cortical collecting duct (CD) cystogenesis, but worsening proximal tubular and glomerular dilations, without affecting kidney function, cilia length, inflammation, and ERK, STAT3, and O-GlcNAc signaling. In a late-onset, adult ADPKD mouse model, deletion of *Thm1* improves kidney function and reduces cystogenesis, inflammation, cilia length, and ERK, STAT3, and O-GlcNAc signaling.

Embryonic or perinatal mutation of *Kif3a, Ift88, Tulp3* or *Thm1* causes renal cystic disease^24, 21, 44, 45^, indicating that all these genes are required for kidney development. Thus, their differential effects in juvenile ADPKD mouse models may reflect differences in their cellular functions, such as cilia length regulation/ciliary homeostasis, ciliary entry or exit of different protein cargo, and regulation of signaling pathways. While IFT-B and -A complexes co-operate in ciliary homeostasis, these complexes also have distinct roles. IFT-B genes mediate anterograde IFT, while IFT-A genes mediate retrograde IFT as well as ciliary entry of membrane and signaling proteins. In contrast, *Tulp3* does not mediate IFT, but is required for ciliary entry of certain membrane-associated and signaling molecules^43^. Additionally, IFT-B and -A mutants can have opposing signaling phenotypes. Any of these functions can be explored to account for the differing juvenile dko phenotypes.

Increased cilia lengths on renal epithelia of several ADPKD mouse models, *PKD1*^*RC/RC*^, *Pkd1* and *Pkd2* cko mice^46, 47^, and recently, on human ADPKD tissue^22^, have been reported. Our data substantiate that polycystin dysfunction causes increased cilia length in mouse models as well as in human ADPKD, suggesting ciliary mechanisms are likely conserved between mouse and human. Further, our data show that in addition to genotype, cilia structure varies by renal tubule segment and maturation, suggesting that factors within a tubule’s microenvironment affect cilia length. Indeed, multiple factors including intracellular Ca^2+^ and cAMP, oxidative stress, cytokines, and fluid flow, affect cilia length of renal epithelial cells^48–50^, indicating that cilia length may be finely regulated in order to maintain renal tubular structure and function. In support of an ameliorative effect of reduced cilia length in ADPKD, inhibition of cilia disassembly in *Pkd1* cko mice increased renal cilia length and exacerbated ADPKD^51^, while in the *jck* non-orthologous PKD mouse model, which also has increased renal cilia lengths, pharmacological shortening of primary cilia was associated with attenuated PKD^52^.

Reduced *Ccl2* and Wnt signaling in *Pkd; Ift-B* dko mice^20, 22^, and reduced P53 enhancing cilia disassembly in *Pkd; Ift-B* dko cells^42^ are potential mechanisms by which ablation or shortening of primary cilia attenuate ADPKD. In addition to detecting ligands, primary cilia also detect mechanical cues. Although mechanosensing by primary cilia and the polycystins has been controversial, recent studies have renewed interest in a potential mechanosensory role for the polycystins, particularly regarding tissue microenvironment stiffness^53, 54^. If sensing of physical forces in the tissue microenvironment is essential to maintaining renal tubular function, then other mechanical cues that would change with cyst growth include shear stress and intraluminal pressure. Cilia length itself could then also be a possible contributing factor in PKD severity. Supporting a role for cilia length, a recent study has shown that primary cilia of proximal tubule epithelial cells transduce shear stress into metabolic pathways that culminate in oxidative phosphorylation^55^. Finally, by extrapolating findings of cilia studies from the cancer field^56^, cilia of not only renal tubular epithelial cells, but of interstitial cells might also affect signaling and disease severity.

Our data are the first to reveal increased O-GlcNAcylation in cyst-lining renal epithelial cells of both juvenile and adult ADPKD mouse models, suggesting increased O-GlcNAcylation may be a feature of ADPKD. Increased O-GlcNAcylation is a pathologic feature of diabetic nephropathy^57, 58^, and in rodent models, has shown to promote various aspects of chronic kidney disease^59, 60^ and also renal fibrosis^61^. In contrast, in a mouse model of contrast-induced acute kidney disease, an acute increase in O-GlcNAcylation was protective^62^, emphasizing important differences between chronic and acute increases in O-GlcNAcylation. While acute changes are adaptive and necessary to maintain cellular health and metabolism, chronic changes are likely to contribute to pathology^63^. Chronic elevation of O-GlcNAcylation in cancer cells promotes tumor growth, and cancer cells show the Warburg effect, suggesting similar cellular metabolic alterations between ADPKD and cancer. A recent study has demonstrated that O-GlcNAcylation of Phosphoglycerate kinase 1 (PGK1), which catalyzes the first ATP molecule in glycolysis, activates PGK1 to enhance lactate production and reduce mitochondrial oxidative phosphorylation, promoting the Warburg effect^39^. Perhaps a similar mechanism may occur in ADPKD cells.

OGT localizes to the pericentriolar region during the early phases of ciliogenesis^56^, and perturbation of O-GlcNAcylation affects cilia length^40, 41^. Further, the ciliary structural defects caused by *Ogt* deletion suggest impaired centriole formation and IFT^56^. Chronic hyper-O-GlcNAcylation in diabetic tissues results in ciliary defects^64^, demonstrating a causative link between misregulation of O-GlcNAcylation and defective ciliary homeostasis in a disease context. Since increased renal cilia lengths are associated with ADPKD renal cystogenesis^46, 47^, the regulation of O-GlcNAcylation on ciliogenesis could be another mechanism by which altered O-GlcNAcylation can affect ADPKD. Elucidating the mechanisms by which O-GlcNAc is upregulated in ADPKD and alters cellular metabolism and ciliogenesis could reveal novel mechanisms and therapeutic targets.

In summary, our data demonstrate for the first time the role of IFT-A deficiency in an ADPKD context, revealing differential effects between nephron segments and between developing and mature renal microenvironments. Our findings also reveal for the first time that O-GlcNAcylation is increased in ADPKD. We propose that as a regulator of ciliary homeostasis and of the balance between glycolysis and oxidative phosphorylation, increased O-GlcNAc may drive certain key ADPKD pathological processes.

## Materials and Methods

### Generation of mice

*Pkd1*^*flox/flox*^, *Pkd2*^*flox/flox*^ and *ROSA26-Cre* mice were obtained from the Jackson Laboratories (Stock numbers 010671, 017292 and 004847, respectively). Generation of *Thm1* cko mice has been described previously^24^: *Thm1*^*aln/+*^; *ROSA26Cre*^*ERT+*^ male mice were mated to *Thm1*^*flox/flox*^ females. *Pkd1* floxed alleles were introduced into the colony to generate *Thm1*^*flox/flox*^;*Pkd1*^*flox/flox*^ or *Thm1*^*flox/flox*^;*Pkd1*^*flox/+*^ females and *Pkd1*^*flox/flox*^; *Thm1*^*aln/+*^, *ROSA26-Cre*^*ERT/+*^ males, which were mated. Similarly, *Pkd2* floxed alleles were introduced into the colony to generate *Thm1*^*flox/flox*^;*Pkd2*^*flox/flox*^ or *Thm1*^*flox/flox*^;*Pkd2*^*flox/+*^ females and *Pkd2*^*flox/flox*^; *Thm1*^*aln/+*^, *ROSA26-Cre*^*ERT/+*^ males. To generate early-onset *Pkd2* models, *Thm1*^*flox/flox*^;*Pkd2*^*flox/flox*^ or *Thm1*^*flox/flox*^;*Pkd2*^*flox/+*^ nursing mothers mated to *Pkd2*^*flox/flox*^; *Thm1*^*aln/+*^, *ROSA26-Cre*^*ERT/+*^ males were injected intraperitoneally with tamoxifen (8mg/40g; Sigma) at postnatal day 0 (P0) to induce gene deletion. Offspring were analyzed at P21. To generate late-onset *Pkd1* models, offspring from matings between *Thm1*^*flox/flox*^;*Pkd1*^*flox/flox*^ or *Thm1*^*flox/flox*^;*Pkd1*^*flox/+*^ females and *Pkd2*^*flox/flox*^; *Thm1*^*aln/+*^, *ROSA26-Cre*^*ERT/+*^ males were injected intraperitoneally with tamoxifen (8mg/40g) at P35. To generate late-onset *Pkd2* models, offspring from matings between *Thm1*^*flox/flox*^;*Pkd2*^*flox/flox*^ or *Thm1*^*flox/flox*^;*Pkd2*^*flox/+*^ females and *Pkd2*^*flox/flox*^; *Thm1*^*aln/+*^, *ROSA26-Cre*^*ERT/+*^ males were injected intraperitoneally with tamoxifen (8mg/40g) at P28. Mice were analyzed at 6 months of age. All mouse lines were maintained on a pure C57BL6/J background (backcrossed 10 generations). All animal procedures were conducted in accordance with KUMC-IACUC and AAALAC rules and regulations.

### Blood Urea Nitrogen Measurements

Mouse trunk blood was collected in Microvette CB 300 Blood Collection System tubes (Kent Scientific), and centrifuged at 1800g at room temperature for 10 minutes to collect serum. BUN was measured using the QuantiChrom Urea Assay Kit (BioAssay Systems) according to the manufacturer’s protocol.

### Histology

Kidneys were bisected transversely, fixed in 10% formalin for several days, then processed in a tissue processor and embedded in paraffin. Tissue sections (7μm) were obtained with a microtome. Sections were deparaffinized, rehydrated through a series of ethanol washes, stained with hematoxylin and eosin (H&E), and mounted in Permount (ThermoFisher). Images were taken with a Nikon 80i microscope equipped with a Nikon DS-Fi1 camera. Cystic areas of H&E-stained sections were quantified using ImageJ.

### Immunofluorescence

Following deparaffinization and rehydration, tissue sections were subjected to antigen retrieval. Tissue sections were steamed for 15 minutes in Sodium Citrate Buffer (10 mM Sodium Citrate, 0.05% Tween 20, pH 6.0), returned to room temperature, rinsed 10 times in distilled water, washed 5 minutes in PBS, incubated for 5 minutes in 1% SDS in PBS based on a method by Brown et al., 1996^65^, then washed 3 times in PBS. Sections were blocked with 1% BSA in PBS for 1 hour at room temperature, then incubated with primary antibodies alone or together with lectins (Table S1) overnight at 4°C. Sections were washed three times in PBS, then incubated with secondary antibodies (Table S2) for 1 hour. After three washes of PBS, sections were mounted with DAPI Fluoromount-G (Electron Microscopy Sciences). Staining was imaged using a Nikon 80i microscope with a photometrics camera or a Nikon Eclipse TiE attached to an A1R-SHR confocal, with an A1-DU4 detector, and LU4 laser launch.

### Immunohistochemistry

Immunohistochemistry was performed as described^66^ using the following primary and secondary antibodies (Tables S1, S2). Once desired color was obtained, sections were counterstained with haemotoxylin. Staining was imaged using a Zeiss A1 microscope with a Axiocam 105 color camera.

### X-gal staining

Kidneys of P0 mice harboring the *Thm1-lacZ* allele (KOMP Repository) were dissected, fixed for 5 min in 10% formalin, washed in lacZ wash buffer, then stained in X-gal solution at 37°C overnight as described^67^. Kidneys were fixed in 10% formalin at 4°C overnight, processed, embedded in paraffin, and sectioned. Sections were rehydrated and either mounted in Permount or labelled with DBA or LTL lectin and mounted in DAPI Fluoromount-G. Sections were imaged using brightfield and confocal microscopy as described^68^.

### Western blot

Kidney pieces (1/3 of kidney) were homogenized in Passive Lysis Buffer (Promega) containing protease inhibitors (Thermofisher). BCA assays (Pierce, ThermoFisher) were performed according to manufacturer’s instructions. Western blots were performed as described^66^, using primary and secondary antibodies (Tables S1 and S2).

### Scanning electron microscopy

SEM on renal tissue was performed as described^69^. Anesthetized mice were perfused with cold 2% glutaraldehyde in 0.1M cacodylate buffer, pH range 6.8-7.4. Kidney cortices were dissected into small pieces and fixed in 2% glutaraldehyde in 0.1M cacodylate buffer at 4°C overnight, then washed in 0.1 M Na-cacodylate, pH 7.4. Samples were post-fixed with 1% OsO4 in 0.1 M Na-cacodylate buffer for 30 minutes, then dehydrated in an ethanol series, followed by hexamethyldisilazane (HMDS; Electron Microscopy Sciences). Samples were mounted onto metal stubs and sputter coated with gold. Samples were viewed and imaged using a Hitachi S-2700 Scanning Electron Microscope equipped with a Quartz PCI digital capture.

### qPCR

RNA was extracted using Trizol (Life Technologies), then reverse transcribed into cDNA using Quanta Biosciences qScript cDNA mix (VWR International). qPCR was performed in duplicate using Quanta Biosciences Perfecta qPCR Supermix (VWR International) in a BioRad CFX Connect Real-Time PCR Detection System with the following primers (Table S3)^24, 70^.

### ADPKD and normal human kidney sections

ADPKD (K386, K408, K423) and normal human kidney (NHK; K357, K402, K419) sections were obtained from the PKD Biomarkers, Biomaterials, and Cellular Models Core in the Kansas PKD Center. The protocol for the use of discarded human tissues complied with federal regulations and was approved by the Institutional Review Board at KUMC^66^.

### Statistics

GraphPad Prism 8 software was used to perform statistical analyses. For analyses of more than two groups, ANOVA or Kruskal-Wallis tests were used to determine statistical significance (p<0.05) of data with or without a normal distribution, respectively. For analysis of two groups, an unpaired t-test was used.

## Acknowledgements

We thank members of the KUMC Department of Anatomy and Cell Biology and the Jared Grantham Kidney Institute for helpful discussions. We thank Jing Huang of the KUMC Histology Core, which is supported by NIH U54HD090216 and NIH P30GM122731. We also thank Pat St. John and Larysa Stroganova of the KUMC Electron Microscopy Research Laboratory, which is supported by NIH P20GM104936. This work was also supported by a K-INBRE Summer Student Award to JTC (K-INBRE P20GM103418), the PKD Biomaterials and Biomarkers Repository Core in the Kansas PKD Research and Translational Core Center (NIH P30DK106912 to JPC), R01DK108433 to MS, and R01DK103033 to PVT.

## Contributions

WW, LMS, MAK, HHW, TSP, BAA, DTJ, RD, JTC, AC, MTP, MS, DPW, and PVT performed experiments. WW, LMS, MAK, HHW, TSP, BAA, DTJ, RD, JTC, AC, MTP, MS, CS, DPW, JPC and PVT analyzed and interpreted data. WW, LMS, BAA, and PVT designed research. WW, LMS, and PVT wrote the manuscript. All authors revised and approved the final manuscript.

